# Exact solutions of coupled multispecies linear reaction–diffusion equations on a uniformly growing domain

**DOI:** 10.1101/026229

**Authors:** Matthew J Simpson, Jesse A Sharp, Liam C Morrow, Ruth E Baker

## Abstract

Embryonic development involves diffusion and proliferation of cells, as well as diffusion and reaction of molecules, within growing tissues. Mathematical models of these processes often involve reaction–diffusion equations on growing domains that have been primarily studied using approximate numerical solutions. Recently, we have shown how to obtain an exact solution to a single, uncoupled, linear reaction–diffusion equation on a growing domain, 0 *< x < L*(*t*), where *L*(*t*) is the domain length. The present work is an extension of our previous study, and we illustrate how to solve a system of coupled reaction–diffusion equations on a growing domain. This system of equations can be used to study the spatial and temporal distributions of different generations of cells within a population that diffuses and proliferates within a growing tissue. The exact solution is obtained by applying an uncoupling transformation, and the uncoupled equations are solved separately before applying the inverse uncoupling transformation to give the coupled solution. We present several example calculations to illustrate different types of behaviour. The first example calculation corresponds to a situation where the initially–confined population diffuses sufficiently slowly that it is unable to reach the moving boundary at *x* = *L*(*t*). In contrast, the second example calculation corresponds to a situation where the initially–confined population is able to overcome the domain growth and reach the moving boundary at *x* = *L*(*t*). In its basic format, the uncoupling transformation at first appears to be restricted to deal only with the case where each generation of cells has a distinct proliferation rate. However, we also demonstrate how the uncoupling transformation can be used when each generation has the same proliferation rate by evaluating the exact solutions as an appropriate limit.

## Introduction

Several processes during embryonic development are associated with the migration and proliferation of cells within growing tissues. A canonical example of such a process is the development of the enteric nervous system (ENS) [1–5]. This involves a population of precursor cells that is initially confined towards the oral end of the developing gut tissue. Cells within the population undergo individual migration and proliferation events, leading to a population–level front of cells that moves toward the anal end of the gut [6]. The spatial distribution of the population of cells is also affected by the growth of the underlying gut tissue [7, 8]. Normal development of the ENS requires that the moving front reaches the anal end of the developing tissue. Conversely, abnormal ENS development is thought to be associated with situations where the front of cells fails to reach the anal end of the tissue [6, 7].

Previous mathematical models of ENS development involve reaction–diffusion equations on a growing domain [6, 9]. These partial differential equation models have been solved numerically, and the numerical solutions used to investigate the interaction between the rates of cell migration, cell proliferation and tissue growth. The interaction between these processes is of interest as it has been shown that altering the relative rates of cell migration, cell proliferation and tissue growth has an important impact on whether the moving cell front can overcome the effects of tissue growth and completely colonize the growing tissue [6, 9]. Previous analysis of these types of models has shown that successful colonization requires that: (i) there is a sufficiently large number of cells present at *t* = 0; (ii) the migration rate of cells is sufficiently large; (iii) the proliferation rate of cells is sufficiently large; and (iv) the rate of growth of the underlying tissue is sufficiently small [6, 9].

All initial studies examining the solution of reaction–diffusion equations on growing domains focused on interpreting numerical solutions of the governing equations [6, 9–19]. More recently, we have shown how to obtain an exact analytical solution of a single species, uncoupled, linear reaction–diffusion equation on a growing domain [20, 21]. The aim of the present study is to extend our previous analysis by presenting a framework that can be used to construct the exact solution of a system of coupled, multispecies, linear reaction–diffusion equations on a growing domain. This means that in the present study we consider a system of coupled partial differential equations on a growing domain, and our approach is relevant to an arbitrary number of coupled partial differential equations. The model we analyze can be used to study the spatial and temporal distributions of different generation of cells within a motile and proliferative cell population on a growing domain. To motivate our model, Figure 1(a) illustrates a cell lineage tree for a birth process in which the different generations are identified. Traditional applications of reaction–diffusion models make no distinction between cells of different generations [22–25] whereas more recent analysis has sought to make a distinction between different generations on a nongrowing domain [26]. The recent work by Cheeseman et al. [26] is novel because it involves re–formulating a standard reaction–diffusion model of cell migration and cell proliferation with the aim of studying the spatial and temporal distribution of different generations of cells on a nongrowing domain. In the present study we use a system of coupled linear reaction–diffusion equations to model the spatial and temporal distribution of each generation on a growing domain. We denote the cell density of the *i*^th^ generation as *C_i_*(*x, t*) for *i* = 1, 2, 3*, …*, and our aim is to find exact solutions of the coupled model. This work is novel since exact solutions of coupled multispecies linear reaction–diffusion equations on a growing domain have not been presented previously.

**Figure 1:**
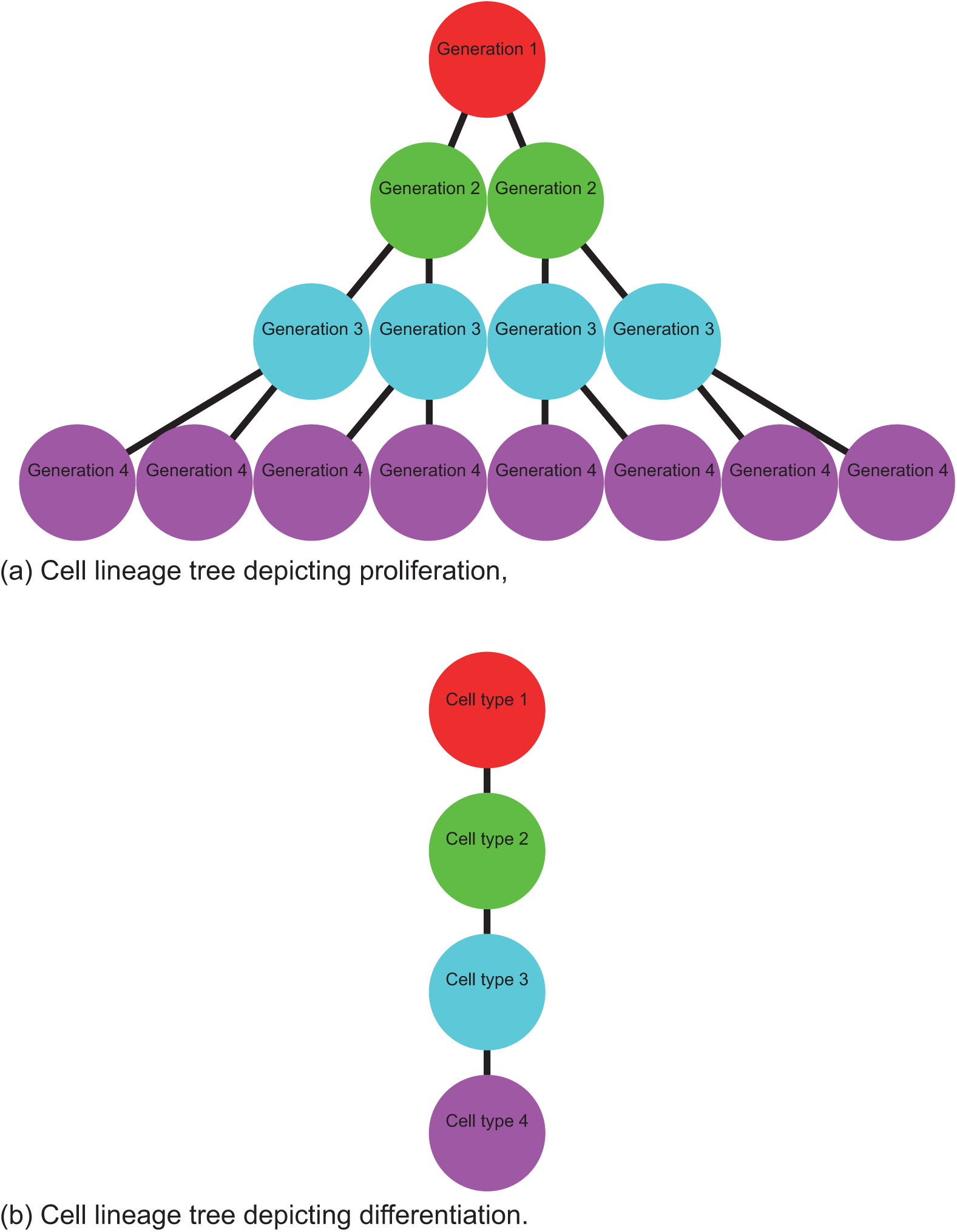
Schematic illustration of two different lineage trees. (a) Lineage tree for a cell proliferation process where each cell gives rise to two daughter cells in the following generation. (b) Lineage tree for a cell differentiation process where each cell undergoes a differentiation process to produce a single cell of a different type.

This manuscript is organized in the following way. First, we outline the mathematical model and the solution strategy. Using the proposed solution method we solve an example problem and present graphical results illustrating some key features of the model, and we always compare the exact solutions with numerical approximations. Although our solution strategy is naturally suited to the most general case where the rate of proliferation of each generation is distinct, we also demonstrate how our approach applies to some special cases in which some of the generations have identical proliferation rates. Additional results relating to the choice of truncation are also presented. Finally, we conclude by summarizing the key findings of our work, and we discuss some other applications for which our analysis is relevant.

## Analysis

We begin by presenting a mathematical model describing the diffusion of a population of cells on a growing domain, where the cells undergo a proliferation process that is depicted schematically in Figure 1(a). This proliferation process means cells in the *i*^th^ generation proliferate to form twice the number of cells in the (*i* + 1)^st^ generation. Assuming that each generation undergoes diffusive movement on a growing domain, we describe the spatial and temporal evolution of the cell density profiles, for each generation, using the following system of coupled linear partial differential equations,

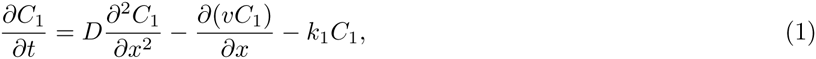

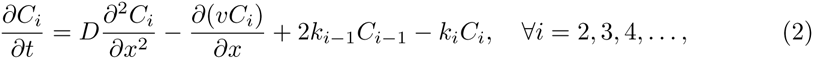

on 0 *< x < L*(*t*). Here, *D* is the cell diffusivity, *v* is the advection velocity associated with domain growth, and *k_i_* is the rate at which cells from the *i*^th^ generation proliferate to produce cells in the next, (*i* + 1)^st^, generation. Note that the factor of two in the production term for generation *i* ≥ 2 reflects the fact that cells from the *i*^th^ generation proliferate to produce twice the number of cells in the (*i* + 1)^st^ generation, as depicted in Figure 1(a).

Our strategy for solving Equations (1)–(2) is valid for a range of initial conditions and boundary conditions. Regardless of the choice of boundary conditions and initial conditions, to solve Equations (1)–(2) we apply Sun and Clement’s uncoupling transformation [27–33], which can be written as

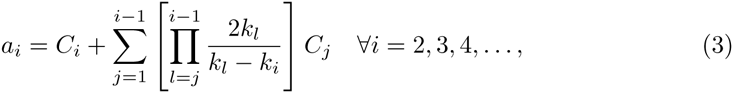

where, for the moment, we require that we have distinct proliferation rates to avoid any singularity in the definition of *a_i_*(*x, t*). Later we will explain how to relax this assumption. Applying the Sun and Clement transformation to Equations (1)–(2) leads to a system of uncoupled partial differential equations,

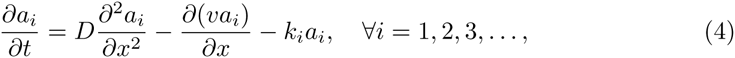

on 0 *< x < L*(*t*), which, at this point, can be solved by using the methods outlined in our previous work for single uncoupled reaction–diffusion equations on growing domains [20, 21]. We note that the solution of Equation (4) can be unbounded when *k_i_* < 0. While we do not outline the entire details of the solution strategy, we will briefly recall the salient features of how to solve Equation (4).

### Domain growth

Domain growth is associated with a velocity field which causes a point at location *x* to move to *x* + *v*(*x, t*)*τ* during a small time interval duration *τ*. We can relate *v*(*x, t*) and *L*(*t*) by considering the expansion of an element of initial width Δ*x* [6],

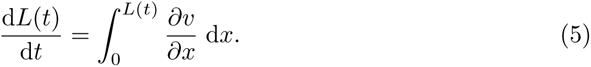

We consider uniform growth conditions where *∂v/∂x* is independent of position, but could depend on time, so that we have *∂v/∂x* = *σ*(*t*) [6, 9–16]. Combining this with Equation (5) gives:

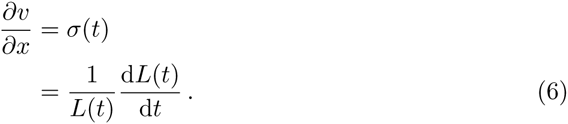

Like previous studies [6, 9, 14], we assume that the domain elongates in the positive *x*–direction with the origin fixed, giving *v*(0*, t*) = 0. Integrating Equation (6) gives

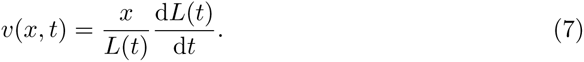

This framework allows us to specify *L*(*t*), for example, by using experimental observations [8], and to use Equation (7) to find the velocity, *v*(*x, t*). For example, exponential growth, *L*(*t*) = *L*(0)e^*αt*^, corresponds to *σ*(*t*) = *α* and *v*(*x, t*) = *αx*. Alternatively, linear growth, *L*(*t*) = *L*(0) + *βt*, corresponds to *σ*(*t*) = *β/*(*L*(0) + *βt*) and *v*(*x, t*) = *xβ/*(*L*(0) + *βt*) [20, 21].

### Solution strategy

To solve Equation (4) we use a Lagrangian mapping, which in this context is also known as a boundary fixing transformation, *ξ* = *x/L*(*t*), giving

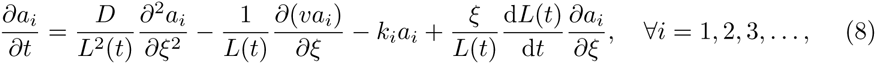

on the fixed domain 0 *< ξ* < 1. Since *v* = *ξ*d*L*(*t*)*/*d*t*, we have

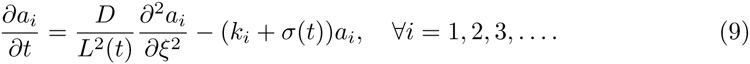

The net reaction term in Equation (9) is the sum of two terms that represent two distinct processes. The first reaction term, *-k*_*i*_*a*_*i*_, is a sink term that is proportional to the rate at which the *i*^th^ generation proliferates to form the (*i* + 1)^st^ generation. The second reaction term, *-σ*(*t*)*a*_*i*_, is proportional to *∂v/∂x*, and since *∂v/∂x >* 0 this is a sink term that represents a dilution effect caused by the domain growth. To simplify Equation (9) we re–scale the time variable, 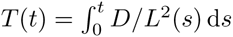[20], so that the coefficient of the diffusive term is constant. This gives

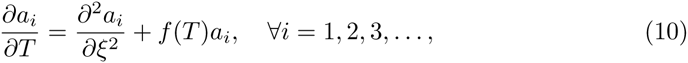

where *f* (*T*) = *-L*^2^(*T*)(*k*_*i*_ + *σ*(*T*))*/D*. Equation (10) can be solved using separation of variables. With zero diffusive flux conditions at both boundaries we have

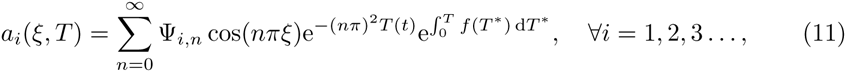

where we choose the Fourier coefficients, ψ_*i*,*n*_, so that *a*_*i*_(*ξ, T*) matches the appropriate initial condition for each component, *i* = 1, 2, 3*, …*. Once the Fourier coefficients have been defined, the exact solution for each uncoupled component can be rewritten in terms of the physical coordinate system, *a*_*i*_(*x, t*), and then re–expressed in terms of the original coupled variables to give *C*_*i*_(*x, t*) for *i* = 1, 2, 3*, …*.

At this point it is worthwhile pointing out how different boundary conditions and initial conditions can be applied. Different initial conditions can be implemented simply by choosing different Fourier coefficients [34]. Applying homogeneous or nonhomogeneous Dirichlet boundary conditions can be implemented by choosing appropriate eigenfunctions in Equation (11) so that the solution satisfies those boundary conditions [21]. The specific examples that we present here in the Results section illustrate how homogeneous Neumann (zero flux) boundary conditions are applied. We choose to focus our examples on using homogeneous Neumann boundary conditions because previous studies have also used similar boundary conditions [6, 20]. We note, however, that greater care is required when applying nonhomogeneous Neumann (non–zero flux) boundary conditions (Supporting Information 1).

## Results

### Distinct reaction rates

Our approach for solving coupled linear reaction–diffusion equations on uniformly growing domains is sufficiently general that it applies to: (i) various types of domain growth functions, *L*(*t*) [20, 21]; (ii) an arbitrary number of generations in the lineage tree [27, 28]; and (iii) arbitrary initial conditions. To demonstrate how our approach applies to a particular problem we will present a suite of results focusing on exponential domain growth, *L*(*t*) = *L*(0)e^*αt*^ with *α >* 0, and, for simplicity, we keep track of the first four generations only by solving

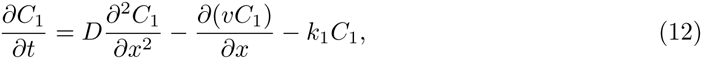

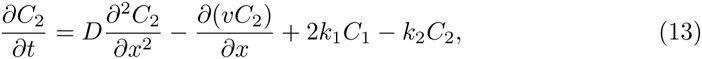

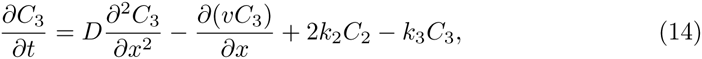

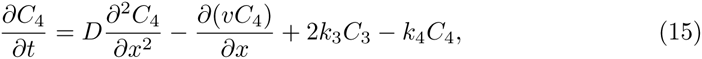

on 0 *< x < L*(*t*). Although all the main results in this work are presented for four generations only, our solution strategy can be adapted to deal with more generations by extending this example in an obvious way. Setting *k*_4_ *>* 0 in this example implies that *C*_4_(*x, t*) will always decay to zero in the long time limit since we have truncated the number of generations to four and we do not explicitly consider the role of the fifth generation. One way of dealing with this is to set *k*_4_ = 0 in the example calculations so that the fourth generation do not proliferate. Another way of dealing with this is to increase the number of generations by including partial differential equation models for *C*_5_(*x, t*), *C*_6_(*x, t*), and so on. However, since this is the first time that these results have been presented we chose to truncate the system after just four generations since we wish to present the results as clearly as possible by working with a modest number of generations. Motivated by Landman’s previous numerical study of ENS development [6], we consider the initial condition

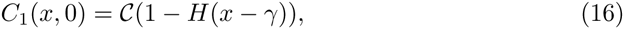

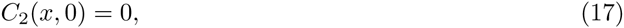

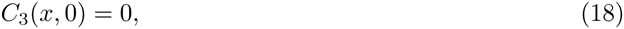

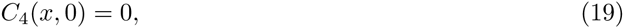

where *H* is the Heaviside function. This initial conditions states that we have some region of the domain, 0 *< x < γ*, initially uniformly occupied by the first generation at density *C*. The remaining portion of the domain, *γ < x < L*(0), is free from cells of the first generation. All other generations are absent at *t* = 0. We apply the Sun and Clement transformation [27, 28], which in this case, can be written as

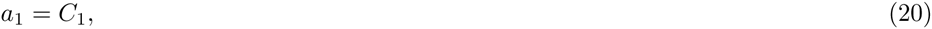

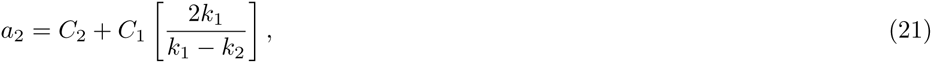

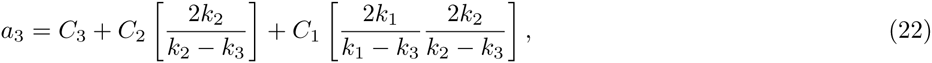

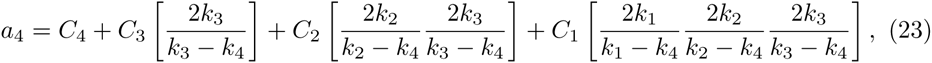

to give four uncoupled partial differential equations. Assuming we have zero diffusive flux boundary conditions at both boundaries, the solutions of the uncoupled partial differential equations can be written as

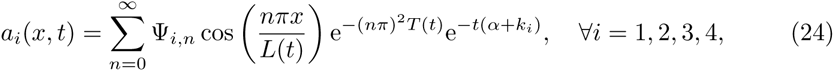

where *L*(*t*) = *L*(0)e^*αt*^ and *T* (*t*) = *D*(1 − e^-2^*αt*)*/*(2^*α*^*L*^2^(0)) [20, 21]. To ensure that *a*_*i*_(*x,* 0) matches the appropriate initial condition, we require

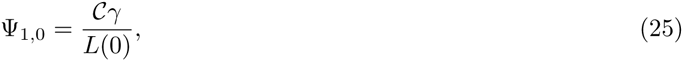

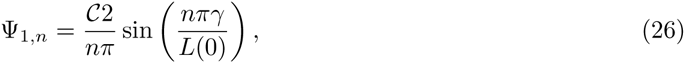

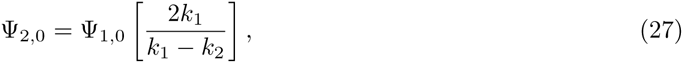

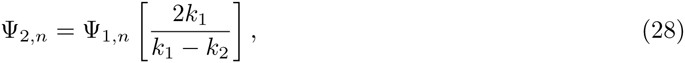

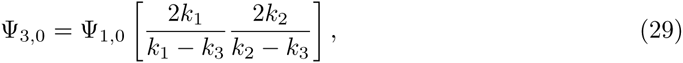

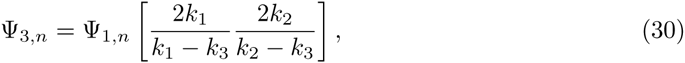

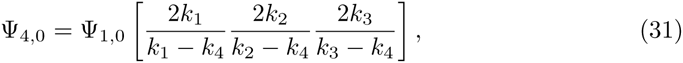

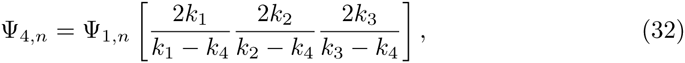

where *n* ∊ ℕ^+^. Given the solutions in the uncoupled format, *a*_*i*_(*x*, *t*), *i* = 1,2,3,4 we then obtain the coupled solutions, *C*_*i*_(*x*, *t*), *i* = 1,2,3,4 using Equations (20)–(23).

Results in Figure 2 show the solutions of Equations (12)–(15) in the case where we have distinct proliferation rates, *k*_1_ ≠ *k*_2_ ≠ *k*_3_ ≠ *k*_4_. The first row shows the initial condition, given by Equations (16)–(19), while the second and third rows show the spatial distribution of each generation and the total density, 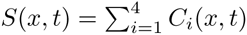, at *t* = 10 and *t* = 20, respectively. Each subfigure contains a plot of the exact solution, truncated after 1000 terms, superimposed on a plot of the numerical solution (Supporting Information 1), and we see that the numerical and exact solutions are visually indistinguishable. Comparing the solutions in Figure 2(a),(f) and (k) indicates that the initial condition is entirely composed of the first generation, whereas by *t* = 20 the first generation is almost absent due to proliferation. In contrast, comparing the results in Figure 2(d), (i) and (n) shows that, initially, the fourth generation is absent and that by *t* = 20 there is a significant population of the fourth generation present on the growing domain. The temporal evolution of the total density, shown in Figure 2(e), (j) and (o), confirms that the spreading cell density profile fails to reach the moving boundary by *t* = 20 [20]. In particular, our exact results indicate that we have *S*(*L*(20), 20) = 0.0000 (correct to four decimal places). Furthermore, if we evaluate the solutions for larger values of *t* we observe that, for this combination of parameters, domain growth dominates and, in effect, the spreading density profile never reaches the moving boundary at *x* = *L*(*t*), and we have *S*(*L*(*t*)*, t*) ≈ 0 [20]. Previous numerical studies of ENS development have pointed out that this kind of result, where the spreading cell density profile fails to reach the end of the growing domain, is consistent with abnormal ENS development [6].

**Figure 2:**
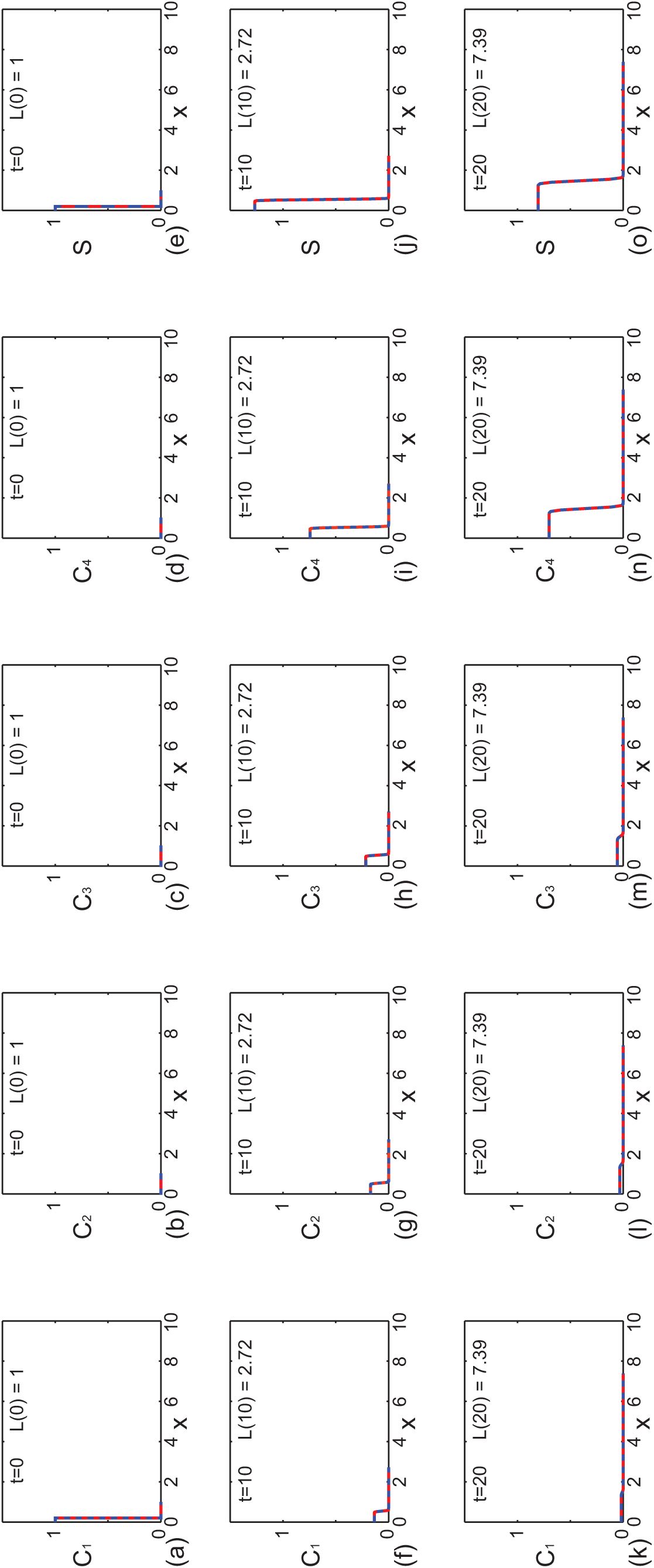
Comparison of exact and numerical solutions of Equations (12)–(15) with distinct reaction rates where colonization fails to occur by. *t* = 20. Profiles in (a)–(e), (f)–(j) and (k)–(o) show *C*_1_(*x, t*), *C*_2_(*x, t*), *C*_3_(*x, t*), *C*_4_(*x, t*) and *S*(*x, t*) at *t* = 0, 10, and 20, respectively. Each subfigure shows the exact solution (solid red) superimposed on the numerical solution (dashed blue). This example corresponds to exponential domain growth with *L*(0) = 1, *L*(10) = e ≈ 2.78 and *L*(20) = e^2^ ≈ 7.39, as indicated in each subfigure. The exact solutions are obtained by truncating the infinite series after 1000 terms and the numerical solutions (Supporting Information 1) correspond to *φξ* = *φt* = 1 *×* 10^−3^. Other parameters are *L*(0) = 1, *a* = 0.1, *C* = 1, *γ* = 0.2, *D* = 1 *×* 10^−5^, *k*_1_ = 0.1, *k*_2_ = 0.2, *k*_3_ = 0.3 and *k*_4_ = 0.

We also present a second set of results, in Figure 3, that are the same as those in Figure 2 with the exception that the diffusivity is increased. Similar to the results in Figure 2 we see that the numerical and exact solutions are visually indistinguishable, and that the density profile of the first generation is present at *t* = 0 and *t* = 10, but is almost absent by *t* = 20. Similarly, the density profile of the fourth generation is identically zero at *t* = 0 but the effects of proliferation mean that the fourth generation is present, and dominates the total population, by *t* = 20. If we compare the evolution of the total density profile, shown in Figure 3(e), (j) and (o), with the evolution of the total density profile in the previous example with smaller *D*, shown in Figure 2(e), (j) and (o), we see that the effect of increasing the diffusivity is that the spreading cell density profile is able to overcome domain growth and colonize the domain. In particular, the exact solutions give *S*(*L*(20), 20) = 0.0085 (correct to four decimal places), which could be interpreted as indicating that the spreading cell density profile has reached the moving boundary at *x* = *L*(*t*) by *t* = 20 [20]. Previous numerical studies of ENS development have pointed out that this kind of result, where the spreading cell population reaches the end of the growing domain, is consistent with normal ENS development [6].

**Figure 3:**
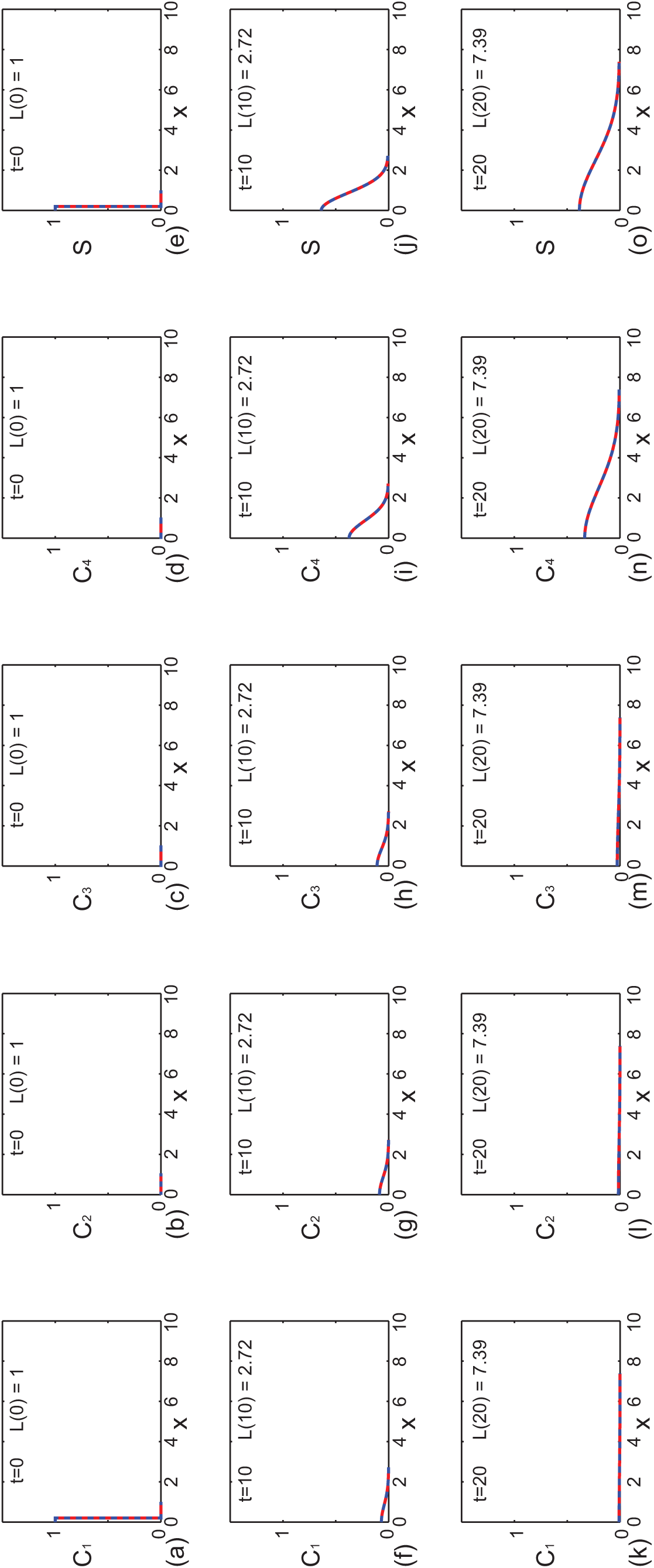
Comparison of exact and numerical solutions of Equations (12)–(15) with distinct reaction rates where colonization occurs occur by. *t* = 20. Profiles in (a)–(e), (f)–(j) and (k)–(o) show *C*_1_(*x, t*), *C*_2_(*x, t*), *C*_3_(*x, t*), *C*_4_(*x, t*) and *S*(*x, t*) at *t* = 0, 10, and 20, respectively. Each subfigure shows the exact solution (solid red) superimposed on the numerical solution (dashed blue). This example corresponds to exponential domain growth with *L*(0) = 1, *L*(10) = e ≈ 2.78 and *L*(20) = e^2^ ≈ 7.39, as indicated in each subfigure. The exact solutions are obtained by truncating the infinite series after 1000 terms and the numerical solutions (Supporting Information 1) correspond to *φξ* = *φt* = 1 *×* 10^−3^. Here we have *L*(0) = 1, *α* = 0.1, *C* = 1, *γ* = 0.2, *D* = 1 *×* 10^−2^, *k*_1_ = 0.1, *k*_2_ = 0.2, *k*_3_ = 0.3 and *k*_4_ = 0.

All exact solutions presented in Figures 2 and 3 are generated using Maple worksheets (Supporting Information 2–3). For all results presented we conservatively truncate the infinite series by retaining the first 1000 terms. Using this approach we find that the computational time required to generate the exact solutions is just a few seconds on a single desktop processor. The numerical solutions of the systems of coupled partial differential equations are generated using code written in FORTRAN 77 [35], and we find that the numerical solutions also requires just a few seconds of computational time on a single desktop processor. Therefore, in summary, there is no particular advantage in terms of computational time requirements to evaluate either the exact or numerical solutions for these problems.

### Repeated reaction rates

As we pointed out in the Introduction, an apparent limitation of the Sun and Clement transformation is that it appears to require distinct proliferation rates to avoid any singularities [27, 28]. We will now show, by example, that it is straightforward to deal with this apparent complication. In particular, we will explain how to obtain exact solutions to Equations (12)–(15) with identical proliferation rates, *k*_1_ = *k*_2_ = *k*_3_ = *k*_4_. The potential issue in solving Equations (12)–(15) with equal proliferation rates is illustrated by visually inspecting the exact solution for *C*_2_,

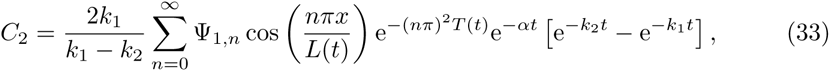

which is indeterminate when *k*_1_ = *k*_2_. This issue can be resolved by evaluating *C*_2_ in the limit as *k*_2_ → *k*_1_ using L’Hopital’s rule, which gives

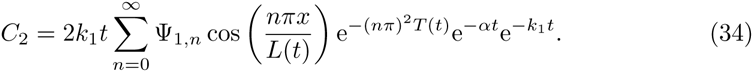

Applying the same approach to the solution of Equations (12)–(15) with *k*^1^ = *k*^2^ = *k*^3^ = *k*^4^ gives,

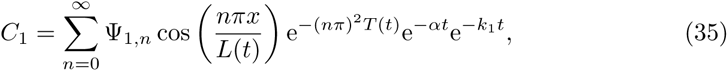

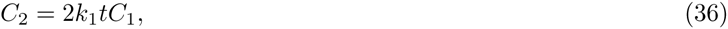

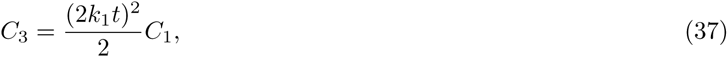

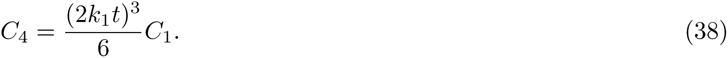

Results in Figure 4 show the solutions of Equations (12)–(15) with *k*_1_ = *k*_2_ = *k*_3_ = *k*_4_. We acknowledge that setting *k*_1_ = *k*_2_ = *k*_3_ = *k*_4_ *>* 0 in Equations (12)–(15) is not biologically realistic since it implies that lim_*t→∞*_ *S*(*x, t*) *≡* 0. However, this exercise of *k*_1_ = *k*_2_ = *k*_3_ = *k*_4_ is mathematically insightful since we wish to illustrate that our general framework for solving the coupled systems of reaction–diffusion equations on a growing domain also applies when we have repeated proliferation rates. The results in Figure 4 are presented in exactly the same format as those in Figures 2 and 3 except that the proliferation rates are equal. As in Figures 2 and 3, the results in Figure 4 indicate that the numerical and exact solutions are visually indistinguishable.

**Figure 4:**
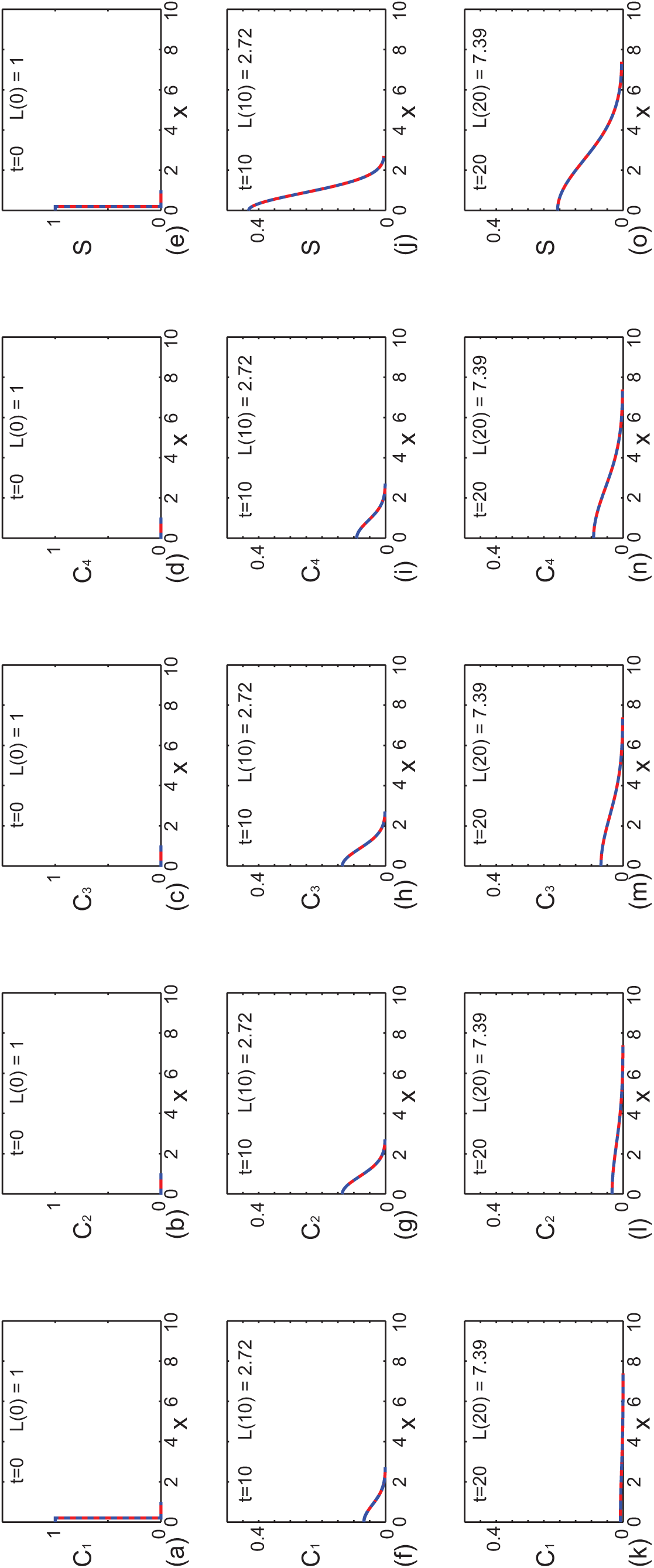
Comparison of exact and numerical solutions of Equations (12)–(15) with equal reaction rates. Profiles in (a)–(e), (f)–(j) and (k)–(o) show *C*_1_(*x, t*), *C*_2_(*x, t*), *C*_3_(*x, t*), *C*_4_(*x, t*) and *S*(*x, t*) at *t* = 0, 10, and 20, respectively. Each subfigure shows the exact solution (solid red) superimposed on the numerical solution (dashed blue). This example corresponds to exponential domain growth with *L*(0) = 1, *L*(10) = e ≈ 2.78 and *L*(20) = e^2^ ≈ 7.39, as indicated in each subfigure. The exact solutions are obtained by truncating the infinite series after 1000 terms and the numerical solutions (Supporting Information 1) correspond to *δξ* = *δt* = 1 *×* 10^−3^. Here we have *L*(0) = 1, *α* = 0.1, *C* = 1, *γ* = 0.2, *D* = 1 *×* 10^−2^, *k*_1_ = *k*_2_ = *k*_3_ = *k*_4_ = 0.1

The example presented in Figure 4 is relevant for the special case where all proliferation rates are identical, with *k*_1_ = *k*_2_ = *k*_3_ = *k*_4_. A similar procedure can be used to obtain the exact solutions in cases where some of the proliferation rates are repeated and others are distinct. For example, the solution of Equations (12)–(15), with *k*_1_ = *k*_2_ = *k*_3_ ≠ *k*_4_, can be written as

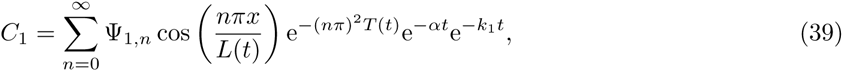

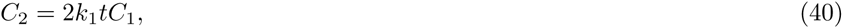

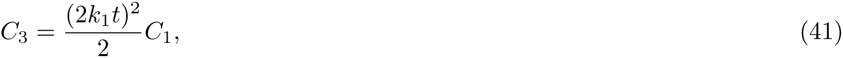

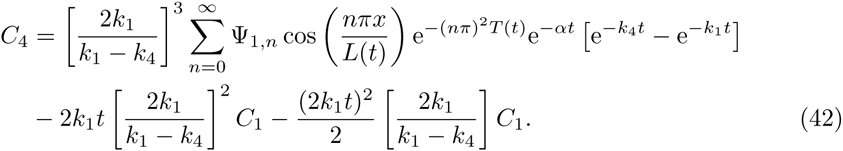

We also compared plots of the numerical solution of Equations (12)–(15), for *k*_1_ = *k*_2_ = *k*_3_ ≠ *k*_4_, with the exact solution, given by Equations (39)–(42), and we observed an excellent match between the exact and numerical solutions (results not shown).

### Choice of truncation

All applications of the solution strategy presented in this work require the infinite series to be truncated after a finite number of terms. For simplicity we always truncate the series very conservatively by retaining the first 1000 terms. The Maple worksheets used to calculate these exact solutions are provided as Supporting Information and these worksheets can be very easily manipulated to explore the effect of varying the level of truncation (Supporting Information 2–3). To demonstrate this, we present additional results in Figure 5(a) showing 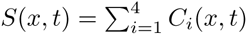 for the same problem considered previously in Figure 3. The profiles in Figure 5(a) compare the exact solution truncated after 1, 2, 5 and 1000 terms. Visual inspection of the profiles indicate that the profile corresponding to 1000 terms is indistinguishable from the profile corresponding to 5 terms. In contrast, the profiles corresponding to 1 and 2 terms in the truncated series are visually distinct. To quantify these trends we plot, in Figure 5(b), *|S*^exact^(*x, t*) − *S*^truncated^(*x, t*)*|*, at *x* = 0 and *t* = 20, where we suppose that the exact solution is given by retaining 1000 terms in the truncated series. Results in Figure 5(b) indicate that truncating after 1000 terms greatly exceeds what is required to ensure that the truncation error is below machine precision since we are unable to distinguish, beyond machine precision, any difference between retaining 10 terms, 100 terms or 1000 terms in the truncated solution. This implies that the truncation error present in Figures 2–4, where we have evaluated the exact solution very conservatively by retaining 1000 terms in the truncated series, is less than machine precision.

**Figure 5:**
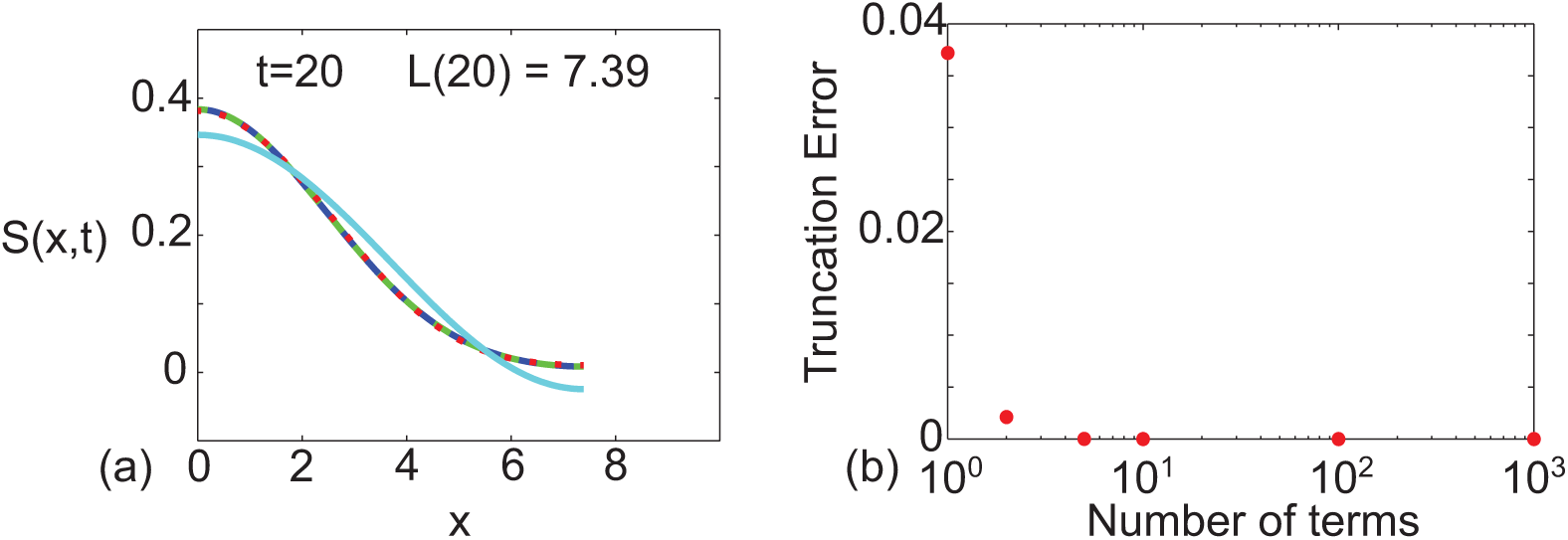
Demonstration of truncation error. Profiles in (a) correspond to the solution of Equations (12)–(15), written in terms of *S*(*x, t*), where 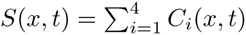. Parameters include *L*(0) = 1, *α* = 0.1, *C* = 1, *σ* = 0.2, *D* = 1 *×* 10^−2^, *k*_1_ = 0.1, *k*_2_ = 0.2, *k*_3_ = 0.3 and *k*_4_ = 0. Results in (a) illustrate the influence of varying the level of truncation in the infinite series by superimposing generated with 1000 terms (solid blue), 5 terms (dashed green), 2 terms (dotted red) and 1 term (solid cyan). Results in (b) show the truncation error, *|S*^exact^(*x, t*) - *S*^truncated^(*x, t*)*|*, at *x* = 0 and *t* = 20, where the exact solution is taken to be the solution generated by truncating after 1000 terms.

Instead of making any prescriptive recommendations about truncating the series, we suggest that any particular application of the solution should involve evaluating the exact solution for the problem of interest iteratively. In each iteration, additional terms in the series should be retained, and the results compared between successive iterations. This process will demonstrate how many terms are required to achieve a desired accuracy. Implementing the exact solution in this way is both straightforward and fast when using the supplied Maple worksheets (Supporting Information 2–3).

## Discussion

In this work we have presented a framework that can be used to calculate the exact solution of a system of coupled linear reaction–diffusion equations on a growing domain. Our work has been motivated by previous numerical studies of ENS development which have used numerical methods to examine the interplay between cell diffusion, cell proliferation and tissue growth in determining whether a cell population, initially confined towards one end of the growing tissue at *x* = 0, can overcome domain growth and reach the other end of the growing tissue at *x* = *L*(*t*) [6, 9]. Most standard models of collective cell spreading make no distinction between different generations of cells [22–25]. In contrast, Cheeseman et al. [26] recently re–formulated a typical reaction–diffusion model of cell migration and cell proliferation so that they could study the spatial and temporal distribution of different generations of cells on a nongrowing domain. Here we use a linear model to make a distinction between different generations of cells in the spreading cell profile and we obtain an exact solution to corresponding system of coupled linear reaction–diffusion equations on a growing domain. Our approach is sufficiently general that it applies to an arbitrary number of generations, an arbitrary initial condition and many choices of the domain growth function, *L*(*t*). This work is novel since we are unaware of any previous work that has presented exact solutions of systems of reaction–diffusion equations on growing domains. However, our approach is limited to dealing with coupled linear reaction–diffusion equations on a one–dimensional growing domain and we suggest that numerical approaches are more appropriate for solving reaction–diffusion equations on two– and three–dimensional growing domains.

While we have motivated our mathematical model by considering a proliferative cell population, our framework can also be adapted to deal with other coupled biological processes on growing domains. For example, the cell lineage tree in Figure 1(b) depicts a cell differentiation process where cells of a particular type differentiate into cells of another type. This kind of cell differentiation process has been incorporated into previous nonlinear coupled multispecies reaction–diffusion models for different types of applications including models of latter stages of ENS development [36, 37] and models of aerosolised skin grafts [38, 39]. If we are interested in applying our technique to solve a linear mathematical model describing cell migration and cell differentiation on a growing domain, we could study a coupled system of linear partial differential equations of the form,

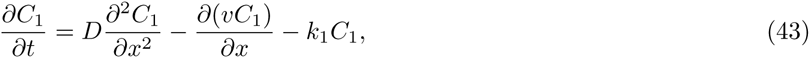

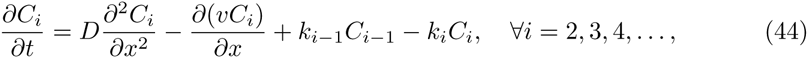

on 0 *< x < L*(*t*). The key difference between Equations (1)–(2) and Equations (43)–(44) is the factor of two in the source terms for *i* ≥ 2. This difference reflects the fact that in the proliferation model cells of each generation proliferate to form twice the number of cells in the next generation whereas cells in the differentiation model differentiate to produce the same number of cells of the next cell type in the cell lineage tree. Applying the Sun and Clement [27, 28] transformation to Equations (43)–(44) proceeds by using a modified version of Equation (4) without the factor of two in the numerator.

The key contribution of our work is to describe a new set of exact mathematical solutions of coupled reaction–diffusion equations on growing domains that have not been presented previously. This contribution is both mathematically and practically relevant because the new exact solutions are motivated by certain problems, such as describing the spatial and temporal distributions of different generations of cells on a growing domain, that cannot be modelled using previous exact solutions [20, 21]. Furthermore, our work is significant because it is the first time, as far as we are aware, that the Sun and Clement transformation [27, 28] has been applied to a problem outside of the porous media literature. Therefore, part of the motivation of this work is to illustrate how the Sun and Clement transformation [27, 28] is relevant to the mathematical biology literature.

Although our comparison of the exact and numerical solutions of Equations (1)–(2) in Figures 2–4 is excellent, our analysis is limited to the study of linear reaction–diffusion equations since we rely on separation of variables and superposition. While many studies of collective cell migration and cell proliferation involve nonlinear partial differential equations [22–25], it is relevant for us consider studying linear partial differential equation models, since they can be viewed as an approximation of nonlinear partial differential equation models. For example, Swanson [40] studied a linearized version of the Fisher-Kolmogorov equation to produce exact analytical solutions that provide insight into the dynamics of tumor spreading. Such linearised models match the solution of the corresponding nonlinear models in the low density limit of the solution which means that the linear model provides a good approximation to the position of the leading edge of the spreading cell population (Supporting Information 1). The fact that the solution of the linear model matches the solution of the nonlinear model at the low density leading edge is both mathematically convenient as well as being of practical interest since many experimental observations of collective cell spreading report results in terms of the position of the low density leading edge of the spreading cell profile [46–48]. We note that similar approximations, which amount to studying nonlinear processes using linearised models, are routinely invoked in many other areas of science and engineering. For example, many nonlinear problems in fluid mechanics [41, 42], civil engineering [43, 44] and chemical engineering [45] are studied, in an approximate sense, by analyzing linearised models. The rationale for studying such linearised models is that they can be solved exactly, thereby providing more general insight than knowledge gathered from repeated numerical simulations.

## Supporting Information Captions

**S1 Supporting Information**. Additional results and discussion.

**S2 Supporting Information**. Maple worksheets to calculate exact solution with distinct proliferation rates.

## References

1. Le Douarin NM, Teillet MA (1973) The migration of neural crest cells to the wall of the digestive tract in avian embryo. J Embryol Exp Morphol. 30: 31–48.

2. Newgreen DF, Erikson CA (1986) The migration of neural crest cells. Int Rev Cytol. 103: 89–145.

3. Gershon MD, Ratcliffe EM (2004) Developmental biology of the enteric nervous system: Pathogenesis of Hirschsprung’s disease and other congenital dysmotilities. Semin Pediatr Surg. 13(4): 224–235.

4. Newgreen DF, Dufour S, Howard MJ, Landman KA (2013) Simple rules for a “simple” nervous system? Molecular and biomathematical approaches to enteric nervous system formation and malformation. Dev Biol. 382: 305–319.

5. Young HM, Bergner AJ, Simpson MJ, McKeown SJ, Hao MM, Anderson CR, Enomoto H (2014) Colonizing while migrating: how do individual enteric neural crest cells behave? BMC Biol. 12: 23.

6. Landman KA, Pettet GJ, Newgreen DF (2003) Mathematical models of cell colonization of uniformly growing domains. Bull Math Biol. 65: 235–262.

7. Newgreen DF, Southwell B, Hartley L, Allan IJ (1996) Migration of enteric neural crest cells in relation to growth of the gut in avian embryos. Acta Anat. 157: 105–115.

8. Binder BJ, Landman KA, Simpson MJ, Mariani M, Newgreen DF (2008) Modeling proliferative tissue growth: A general approach and an avian case study. Phys Rev E. 78: 031912.

9. Simpson MJ, Landman KA, Newgreen DF (2006) Chemotactic and diffusive migration on a nonuniformly growing domain: numerical algorithm development and applications. J Comp Appl Math. 192: 282–300.

10. Crampin EJ, Gaffney EA, Maini PK (1999) Reaction and diffusion on growing domains: scenarios for robust pattern formation. Bull Math Biol. 61: 1093–1120.

11. Crampin EJ, Hackborn WW, Maini PK (2002) Pattern formation in reaction-diffusion models with nonuniform domain growth. Bull Math Biol. 64: 747–769.

12. Kondo S, Asai R (1995) A reaction-diffusion wave on the skin of the marine angelfish pomacanthus. Nature. 376: 765–768.

13. Painter KJ, Maini PK, Othmer HG (1999) Stripe formation in juvenile pomacanthus explained by a generalized Turing mechanism with chemotaxis. Proc Natl Acad Sci USA. 96: 5549–5554.

14. Baker RE, Yates CA, Erban R (2009) From microscopic to macroscopic descriptions of cell migration on growing domains. Bull Math Biol. 72(3): 719–762.

15. Woolley TE, Baker RE, Gaffney EA, Maini PK (2011) Stochastic reaction and diffusion on growing domains: Understanding the breakdown of pattern formation. Phys Rev E. 84: 046216.

16. Yates CA (2014) Discrete and continuous models for tissue growth and shrinkage. J Theor Biol. 350: 37–48.

17. Madzvamuse A, Gaffney EA, Maini PK (2010) Stability analysis of non–autonomous reaction–diffusion systems: the effects of growing domains. J Math Biol. 61: 133–164.

18. Venkataraman C, Lakkis O, Madzvamuse A (2012) Global existence for semilinear reaction–diffusion systems on evolving domains. J Math Biol. 64: 41–67.

19. Dziuk G, Elliott CM (2013) Finite element methods for surface PDEs. Acta Numerica. 22: 289–396.

20. Simpson MJ (2015) Exact solutions of linear reaction-diffusion processes on a uniformly growing domains: Criteria for successful colonization. PLOS ONE. 10(2): e0117949.

21. Simpson MJ, Sharp JA, Baker RE (2015) Survival probability for a diffusive process on a growing domain. Phys Rev E. 91: 042701.

22. Maini PK, McElwain DLS, Leavesley DI (2004) Traveling wave model to interpret a wound-healing cell migration assay for human peritoneal mesothelial cells. Tissue Eng. 10: 475–482.

23. Maini PK, McElwain DLS. Leavesley DI (2004) Travelling waves in a wound healing assay. Appl Math Lett. 17: 575–580.

24. Sherratt JA, Murray JD (1990) Models of epidermal wound healing. Proc R Soc Lond B. 241: 29–36.

25. Sengers BG, Please CP, Oreffo ROC (2007) Experimental characterization and computational modelling of two–dimensional cell spreading for skeletal regeneration. J R Soc Interface. 4: 1107–1117.

26. Cheeseman BL, Newgreen DF, Landman KA (2014) Spatial and temporal dynamics of cell generations within an invasion wave: A link to cell lineage tracing. J Theor Biol. 363: 344–356.

27. Sun Y, Clement TP (1999) A decomposition method for solving coupled multi-species reactive transport equations. Transport Porous Med. 37: 327–346.

28. Sun Y, Petersen JN, Clement TP, Skeen RS (1999) Development of analytical solutions for multispecies transport with serial and parallel reactions. Water Resour Res. 35: 185–190.

29. Sun Y, Lu X, Petersen JN, Buscheck TA (2004) An analytical solution of tetrachloroethylene transport and biodegradation. Transport Porous Med. 55: 301–308.

30. Sun Y, Buscheck TA, Hao Y (2008) Modeling reactive transport using exact solutions for first-order reaction networks. Transport Porous Med. 71: 217–231.

31. Sun Y, Buscheck TA, Hao Y (2012) An analytical method for modeling first-order decay networks. Comput Geosci. 39: 86–97.

32. Srinivasan V, Clement TP (2008) Analytical solutions of sequentially coupled one-dimensional reactive transport problems - Part I: mathematical derivations. Adv Water Resour. 31: 203–218.

33. Srinivasan V, Clement TP (2008) Analytical solutions of sequentially coupled one-dimensional reactive transport problems - Part II: special cases, implementation and testing. Adv Water Resour. 31: 219–232.

34. Haberman R (2004) Applied partial differential equations: with Fourier series and boundary value problems. New York, Prentice Hall.

35. Koffman EB, Friedman FL (1993) FORTRAN with engineering applications. Fifth edition. Reading, Massachusetts, Addison Wesley.

36. Trewenack AJ, Landman KA (2009) A traveling wave model for invasion by precursor and differentiated cells. Bull Math Biol. 71: 291–317.

37. Hou Z (2013) Analysis of a model arising from invasion by precursor and differentiated cells. Int J Diff Equations. 2013: 314173.

38. Denman PK, McElwain DLS, Harkin DG, Upton Z (2007) Mathematical modelling of aerosolised skin grafts incorporating karatinocyte clonal subtypes. Bull Math Biol. 69: 157–179.

39. Denman PK, McElwain DLS, Norbury J (2007) Analysis of travelling waves associated with modelling of aerosolised skin grafts. Bull Math Biol. 69: 495–523.

40. Swanson KR (2008) Quantifying glioma cell growth and invasion in vitro. Math Comput Model. 47: 638–648.

41. Crowley S, Porter R (2012) The effect of slatted screens on waves. J Eng Math. 76: 33–57

42. Simpson MJ, Jazaei F, Clement TP (2013) How long does it take for aquifer recharge or aquifer discharge processes to reach steady state? J Hydroo. 501: 241–248.

43. Bear J (1972) Dynamics of fluids in porous media. New York, American Elsevier Publishing.

44. Haitjema HM (1995) Analytic element modeling of groundwater flow. San Diego, Academic Press.

45. Landman KA, White LR (1997) Predicting filtration time and maxmizing throughput in a pressure filter. AIChE Journal. 43: 3147–3160.

46. Kam Y, Guess C, Estrada L, Weidow B, Quaranta V (2008) A novel circular invasion assay mimics in vivo invasive behaviour of cancer cell lines and distinguishes single–cell motility in vitro. BMC Cancer. 8: 198.

47. Treloar KK, Simpson MJ (2013) Sensitivity of edge detection methods for quantifying cell migration assays. PLOS ONE. 8(6): e67389.

48. Treloar KK, Simpson MJ, McElwain DLS, Baker RE (2014) Are in vitro estimates of cell diffusivity and cell proliferation rate sensitive to assay geometry? J Theor Biol. 356: 71–84.

